# Disturbance of Information in Superior Parietal Lobe during Dual-task Interference in a Simulated Driving Task

**DOI:** 10.1101/2020.07.28.224394

**Authors:** Mojtaba Abbaszadeh, Gholam-Ali Hossein-Zadeh, Shima Seyed-Allaei, Maryam Vaziri-Pashkam

## Abstract

Performing a secondary task while driving causes a decline in driving performance. This phenomenon, called dual-task interference, can have lethal consequences. Previous fMRI studies have looked at the changes in the average brain activity to uncover the neural correlates of dual-task interference. From these results, it is unclear whether the overall modulations in brain activity result from general effects such as task difficulty, attentional modulations, and mental effort or whether it is caused by a change in the responses specific to each condition due to dual-task interference. To overcome this limitation, here, we used multi-voxel pattern analysis (MVPA) to interrogate the change in the information content in multiple brain regions during dual-task interference in simulated driving. Participants performed a lane change task in a simulated driving environment, along with a tone discrimination task with either short or long-time onset difference (Stimulus Onset Asynchrony, SOA) between the two tasks. Behavioral results indicated a robust dual-task effect on driving RT. MVPA revealed regions that carry information about the driving direction, including the superior parietal lobe (SPL), visual, and motor regions. Comparing the decoding accuracies across short and long SOA conditions, we showed lower accuracies in the SPL region in short than long SPA conditions. This change in accuracy was not observed in the visual and motor regions. In addition, the classification accuracy in the SPL was inversely correlated with participants’ reaction time in the driving task. These findings suggest that the dual-task interference in driving may be related to the disturbance of information processing in the SPL region.

**Significance Statement:** During real-world driving, when a driver wants to make a turn at an intersection and simultaneously respond to a cell phone call, his reaction time slows down. This effect is called dual-task interference. Here, we aimed to examine its neural mechanisms using a paradigm that consisted of a driving turn task and a tone discrimination task in a simulated environment. Results showed that the information for the driving turn was disturbed in the superior parietal lobe (SPL) during dual-task interference. We suggest that the driving performance decline in the presence of the secondary task might be related to the disturbance of information in the SPL.

## Introduction

Performing a secondary task during driving causes a decline in driving performance (i.e. (Abbas-Zadeh, Hossein-Zadeh, & Vaziri-Pashkam, 2021; Hibberd, Jamson, & Carsten, 2013; Levy, Pashler, & Boer, 2006). In particular, when the time lag between the onsets of the driving task with the secondary task (hereafter referred to as the Stimulus Onset Asynchrony or SOA) decreases, the reaction times and the accuracies deteriorate. This performance decline as a function of SOA is used as a measure of dual-task interference. Although dual-task interference has been widely studied using behavioral techniques, its underlying neural mechanisms are not well understood. Here, we would like to study the neural correlates of dual-task interference using functional MRI and multi-voxel pattern analysis in a well-controlled simulated driving environment.

Although the dual-task interference affects both tasks in a dual-task condition, the task presented second is more vulnerable to the dual-task effects (Pashler, 1994; Zylberberg, Ouellette, Sigman, & Roelfsema, 2012). Previous studies have suggested several mechanisms for the second-task performance decline in dual-task conditions. Some studies have proposed that these effects are due to the delays in neural processing stages such as sensory processing, response selection, and motor execution (Dux, Ivanoff, Asplund, & Marois, 2006; Dux et al., 2009; Shapiro, Raymond, & Arnell, 1997). Other studies have stated the disruption in the short-term memory encoding as a reason for the second task effects. Furthermore, task switching (Jamadar, Hughes, Fulham, Michie, & Karayanidis, 2010; Kimberg, Aguirre, & D’Esposito, 2000) and task-set reconfiguration (Rogers & Monsell, 1995; Sigman & Dehaene, 2006) have also been reported by previous studies as a reason for dual-task effects.

Numerous dual-task fMRI studies have proposed that dual-task interference arises because the two tasks simultaneously may use the same executive resources in the parietal and frontal cortices (Al-Hashimi, Zanto, & Gazzaley, 2015; Deprez et al., 2013; Hesselmann, Flandin, & Dehaene, 2011; Jiang, 2004; Tombu et al., 2011). These fMRI studies of dual-task interference have used univariate methods for investigating the neural correlates of dual-task interference, looking at either the peak or the time course of the BOLD response. From these results, it is not clear whether the overall modulations in brain activity are a result of general effects such as task difficulty, attentional modulations, and mental effort, or it is caused by a specific change in the neural responses to each condition due to interference in perceptual and cognitive information. A faithful approach to overcome this limitation is investigating the information for each task separately in each brain region using multi-voxel pattern analysis (MVPA) in dual-task conditions (Kamitani & Tong, 2005; Kriegeskorte, Goebel, & Bandettini, 2006). This method provides the opportunity to focus only on specific brain regions that carry information about the driving task and investigate the change in their information content during dual-task interference. Using MVPA and MEG, Marti, King, and Dehaene (2015) investigated the change of the information of two tasks during the dual-task interference in two simple tasks. They showed that the dual-task interference decreases the information of the second task in the parietal cortex between 350-450 ms after stimulus onset. Given the low spatial resolution of MEG, it was impossible to determine the exact brain with the drop of information. Therefore, the combination of MVPA and fMRI can overcome the limitation of their study. To the best of our knowledge, no fMRI study has used multivariate methods to investigate the driving task representation in different dual-task conditions.

Most previous dual-task studies have used two simple tasks to characterize the source of dual-task interference. Based on these studies, it is unclear whether identical neural regions are also associated with performance declines during real-world dual-task conditions, such as performing a secondary task while driving. In the present study, we developed a dual-task experiment in a simulated driving environment to get one step closer to real-world dual-task settings. Although our paradigm does not completely simulate real-world driving, we believe it has some key elements of a lane change scenario in a driving context. Our dual-task paradigm, in which the parameters of previous laboratory dual-task paradigms are still finely controlled, might likely evoke mental states more typical to the ones seen in real-world situations.

A few studies have investigated the brain regions involved in dual-task interference in a simulated driving environment by comparing dual-task with single-task conditions (Al-Hashimi et al., 2015; Just, Keller, & Cynkar, 2008), and they have reported the modulation of activity in the parietal and frontal cortices. One experimental variable that could modulate the amount of dual-task interference and has been overlooked in previous driving dual-task studies is SOA. The manipulation of SOA provides the possibility to localize the regions where their activation correlates with the magnitude of dual-task interference (René Marois & Ivanoff, 2005). Short and long SOA conditions are only different in the timing between the two tasks. Therefore, regions isolated by comparing short and long SOA conditions are potentially more specific than regions isolated by comparing dual- and single-task conditions. We used this approach to isolate regions affected during dual-task interference.

In sum, here, we used a dual-task paradigm in which a driving task was performed concurrently with a tone discrimination task in two dual-task conditions (short SOA and long SOA). Using MVPA, we localized regions that carry information about the driving direction. In these regions, we then investigated if the information for driving direction was disrupted in short compared to the long SOA conditions. Our result showed that the bilateral early visual cortex, right superior parietal lobe (SPL), and right motor cortex have significant information about the driving task. However, the information in the SPL decreased significantly during dual-task interference.

## Material and methods

### Participants

Twenty-four right-handed volunteers (16 females), 20-36 years old with normal or corrected-to-normal vision and no history of neurological or psychiatric disorders participated in the experiment. Participants gave informed consent and received payment for their participation. The ethics committee at the Institute for Research in Fundamental Sciences (IPM) approved the experiment. Four participants were excluded from the analysis due to excessive head movement (> 5 mm) during the scan and the results of 20 participants (12 females) were analyzed.

#### Experimental design and procedure

##### Apparatus

Stimuli were presented on a 32” monitor at the back of the fMRI scanner bore and viewed via a head coil-mounted mirror. Participants responded to the tasks using two fMRI-compatible Current Designs four-button response pads, one for each hand. They responded to the driving task with their left index and middle fingers and the tone task with their right index and middle fingers.

##### Stimuli and Paradigm

The fMRI dual-task paradigm consisted of a driving lane change task and a tone discrimination task. The Unity 3D game engine was used to design the driving environment. The driving environment included a highway with infinite lanes on the two sides, without left/right turns or inclining/declining hills. This was to equalize all trials in terms of visual appearance for the experiment (see Figure 1). The driving stimulus was composed of two rows of traffic cones with three cones in each row (Figure 1). In each trial, traffic cones were unexpectedly displayed on both sides of one of the lanes, and the participants had to steer the vehicle immediately to the lane with the cones and drive through them. The distance between the two rows of cones was such that the vehicle could easily drive through them without collisions. The cones were always presented on the lane immediately to the left or the right of the driving lane so that the participants had to change only one lane per trial. The lane change was performed gradually, and the participants had to hold the corresponding key to direct the vehicle in between the two rows of cones, and then release the key when the vehicle was situated correctly. Any collision with the cones would be registered as an error. The fixation cross was jittered for 100 ms to provide online feedback in case of a collision with the traffic cones.

**Figure 1.**
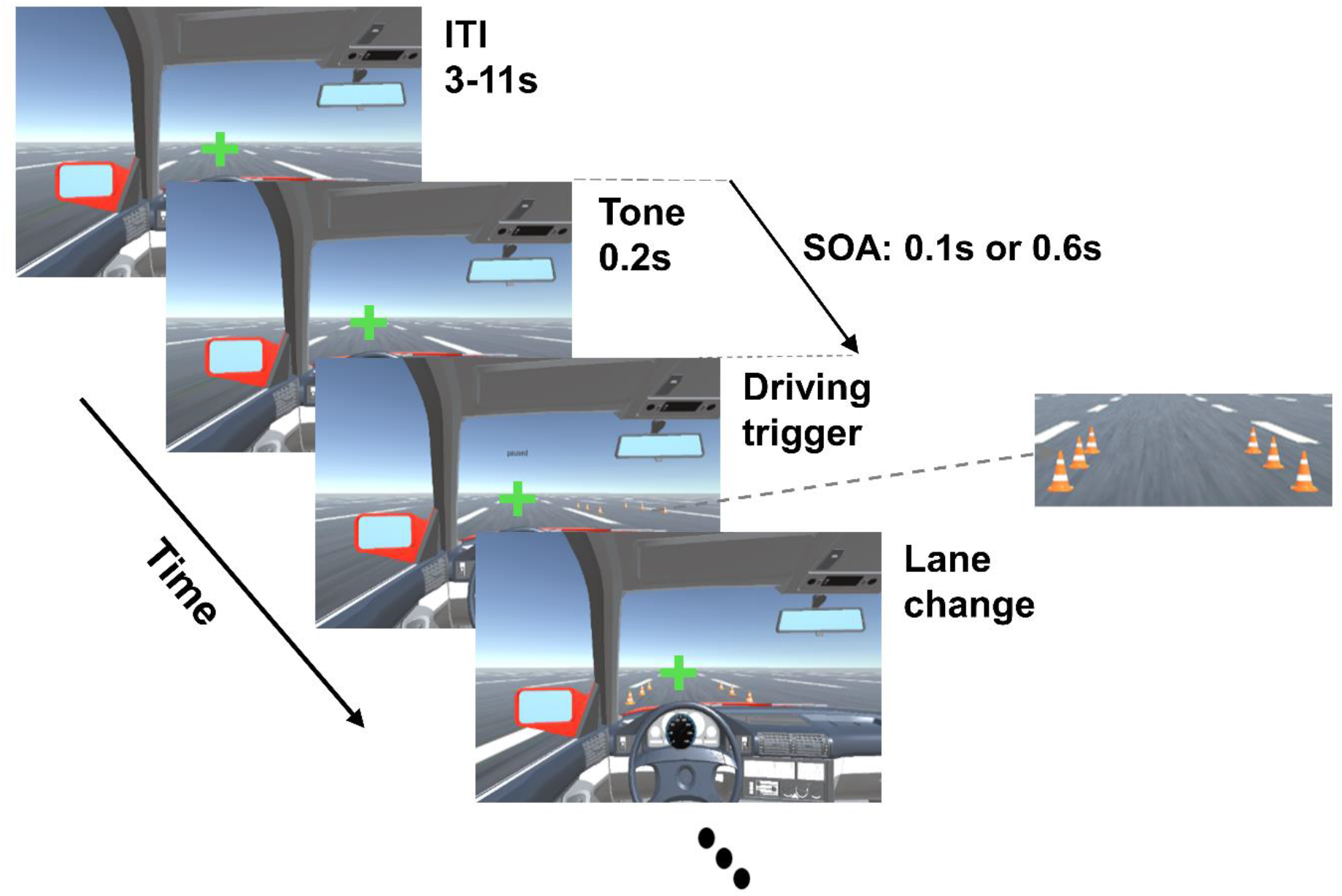
The sequence of events for a sample trial of the dual-task paradigm. The Inter trial interval (ITI) lasted between 3 to 11 s. The tone lasted for 200 ms and the driving trigger was presented 100 or 600 ms after the tone stimulus. Participants had to perform a tone discrimination task immediately after the presentation of the tone and a lane change immediately after the driving stimulus (two rows of cones).

The driving started with a given initial speed which was kept constant. During the experiment, the participant moved to the right or left lanes by pressing the keys using their left hand’s middle and index fingers, respectively. For the tone task, a single pure tone of either a high (800 Hz) or low (400 Hz) frequency was presented for 200 milliseconds. Participants pressed the keys with their right hand’s middle and index fingers to determine whether the tone was a high frequency or a low frequency, respectively. To provide feedback, if participants responded incorrectly, the green fixation cross turned red. The onset of each trial was set to be the presentation time of the tone stimulus and the end of each trial was set to be when the rear end of the car reached the end of the set of traffic cones. Participants were told to focus on the fixation cross at the center of the page and respond as fast as possible to each task that was presented. The performance in the driving task was calculated as the percentage of trials in which the participant passed through the cones without collision. The performance in the tone task was calculated as the number of correct identifications.

The experiment consisted of dual-task and single-task conditions. In the dual-task trials, the two tasks were presented with either short (100 ms) or long (600 ms) SOAs. In the single-task trials, either the driving or the tone task was presented alone. There were eight dual-task conditions: two SOAs x two driving directions (turn right/turn left) x two tone conditions (low/high frequency); and four single-task conditions: two directions x two tone conditions. In the dual-task conditions, the order of the presentation of tasks was fixed so that the tone task was always presented first and the driving task was presented second. Each condition was repeated four times in each run resulting in a total of 48 trials in every run. Each trial lasted 3s with an inter-trial interval varying from 3s to 11s. We used *optseq* software (Dale, 1999) for optimizing the presentation order of trials in each run. The participants completed 12 runs, each lasting 4.8 min (288 sec). Before performing the main experiment, all participants performed two training runs similar to the main experiments. They would proceed to the main experimental runs if their performance was 80% or higher. All participants could reach this threshold.

#### Image acquisition

Structural and functional images were acquired using a Prisma Siemens 3T MRI scanner at the National Brain Mapping Laboratory with a 64-channel head coil for 4 participants and a Tim Trio Siemens 3T MRI scanner with a 32-channel head coil at the IPM Imaging Center for all other participants. The IPM Imaging Center scanner was originally unavailable due to technical reasons, therefore we started the experiments at the National Brain Mapping Laboratory. After the IPM scanner became available we switched to collecting data at IPM. The imaging parameters were kept the same across scanners and the results were similar. T1-weighted images were acquired with gradient echo sequence with 1900 ms repetition time (TR), echo time (TE) = 2.52, the field of view (FOV) = 256 mm, matrix size of 256 x 256, with 192 slices with 1mm thickness, iso voxel size of 1 mm, and flip angle of 9 degrees. These high-resolution images were used for surface reconstruction. Functional images were acquired using a single-shot gradient EPI sequence with TR = 2000 ms, TE = 26 ms, flip angle = 90-degree, matrix size of 64 x 64, FOV = 192 mm, 33 slices with 0.3 mm slice gap, and voxel size of 3 x 3 x 3 mm.

#### Image Preprocessing

Initial image analysis was performed using the Freesurfer image analysis suite, (http://surfer.nmr.mgh.harvard.edu/) and fsfast (Dale, Fischl, & Sereno, 1999). Pattern classification was performed using the CoSMoMVPA toolbox in MATLAB (Oosterhof, Connolly, & Haxby, 2016) and in-house MATLAB code. FMRI preprocessing included 3D motion correction, slice timing correction, and linear and quadratic trend removal. For the whole brain group-level analysis, the anatomical T1-weighted images of each participant were transformed into standardized Freesurfer fsaverage space (Evans et al., 1993). For GLM analysis, data were spatially smoothed with a Gaussian kernel of 6 mm FWHM for univariate analysis, but non-smoothed data were used for multivariate analysis. For every condition, we used a finite impulse response (FIR) basis function (Henson, Rugg, & Friston, 2001) to examine the changes in BOLD response across time. For this purpose, the BOLD response was quantified for 10 post-stimulus time bins with each time bin representing one full-brain volume of 2 s duration. These time bins were time-locked to the onset of the tone stimulus in the dual-task trials and with the onset of the tone stimulus or the driving cones in the single-task tone and driving trials, respectively.

#### Multivariate pattern analysis (MVPA)

A general linear model was first used to estimate beta values for each vertex. We used an FIR analysis to not assume a shape for the hemodynamic response function. We had10 post-stimulus FIR time bins that were time-locked to the onset of the stimulus. For the GLM model, we used these 10 regressors for each of the 12 conditions: eight dual-task conditions (two driving lane change directions x two tone types x two SOAs) and four single-task conditions (two driving lane change direction and two tone types). The focus of the current study was to compare the short and long SOA conditions; therefore, the result of single-task conditions was excluded from the analysis in this paper.

After extracting the beta values, we assessed whether information about the driving direction is encoded differently in the short and long SOA conditions. First, using a searchlight approach (Kriegeskorte et al., 2006) we identified the regions that carried information about the driving direction in either the short SOA or the long SOA conditions. Next, we quantified the differences between the decoding accuracies in the short and long SOA conditions in an ROI analysis.

Using CoSMoMVPA (Oosterhof et al., 2016), we ran a surface-based searchlight analysis (Oosterhof, Wiggett, Diedrichsen, Tipper, & Downing, 2010) on the beta maps of FIR time bin 3 (the peak hemodynamic time-bin) for each participant. The surface-based searchlight was restricted to a mask containing only voxels between the pail and white surfaces of the brain. The pail and white surfaces were generated using Freesurfer. Using the pial and white surfaces an intermediate surface was estimated which was placed between two surfaces. A center vertex on the intermediate surface was selected and then 100 neighboring voxels around the center vertex were chosen based on the geodesic distance as a region. The procedure was performed for all vertices on the intermediate surface that included the entire volume of the gray matter of the brain. The estimated beta value patterns were extracted in every region. Then, in each region, information about the driving direction was assessed by a linear support vector machine classifier. A leave-one-run-out cross-validation procedure was used to evaluate the classification performance (Kamitani & Tong, 2005). The classifier was trained to discriminate between the two classes (turn left vs turn right) from all but one run and tested on the left-out run. This process was repeated for the 12 runs and the resulting performances were averaged to generate the mean classification accuracy for each searchlight center vertex. This analysis was performed separately for the short and long SOA conditions to produce a whole-brain classification accuracy map for each participant and each SOA condition. The average accuracy maps were normalized to a common space (fsaverage). A Gaussian kernel with 6 mm full-width at half maximum was used to smooth the accuracy maps.

To compare the information of the driving task in the short and long SOAs, we performed an ROI analysis using the accuracy maps of the short and long SOA. To define the ROIs, we performed a leave-one-subject-out procedure (Esterman, Tamber-Rosenau, Chiu, & Yantis, 2010) to avoid double-dipping (Kriegeskorte, Simmons, Bellgowan, & Baker, 2009). One participant was left out and by using the remaining participants, a group-level analysis was performed to find clusters that their decoding accuracies were significantly above chance separately for short and long SOAs. Then using a union analysis, we obtained a p-value map equal to the minimum p-value of the short and long SOA. This p-value map was then corrected for multiple comparisons with voxel-wise *p* < 0.001 and cluster thresholded at *p* < 0.05). These clusters related to the 19 participants were selected as ROIs for the left-out 20th participants and the procedure was repeated to obtain independent ROIs for all participants. The mean accuracies across SOAs were compared in these ROIs. To statistically compare the mean accuracy of each ROI for short and long SOAs, we ran a two-way repeated measure ANOVA with SOA (short and long) and FIR time bins as two factors, and p-values were corrected for multiple comparisons with FDR *q* < 0.05 across the ROIs. The comparison of the p-value with the chance level (50%) was performed by a one-sample t-test for each ROI and each time bin. The pairwise comparison of accuracy for the short and long SOA conditions was performed by paired t-test in each ROI and each time bin. The p-values were corrected for multiple comparisons with FDR *q* < 0.05 across time bins for each ROI. We included all trials in the analyses, however, our further exploration of data cleared that removing the error trials did not qualitatively change the results.

## Results

### Behavioral results

We compared the mean RT and accuracy across long and short SOA conditions for the driving and the tone task (Figure 2). For the driving task which was always presented second, mean RT was significantly greater in the short SOA trials than in the long SOA trials (*t*(19) = −13.23, *p* < 0.0001, paired t-test). Although the participants had high accuracy in the driving task (accuracy > 95%), the participants’ accuracy was significantly lower in short compared to long SOA conditions (*t*(19) = −4.06, *p* < 0.001, paired t-test). For the tone task, the RT did not significantly change SOA conditions (*ps* > 0.05), while the accuracy was significantly lower in the short compared to the long SOA trials (*t*(19) = 4.11, *p* < 0.001, paired t-test). These findings show that the performance of the two tasks is influenced by dual-task interference, although the effect is stranger for the reaction time of the driving task (*t*(19) = −4.63, *p* < 0.0001, paired t-test). These findings are consistent with previous studies (Abbas-Zadeh et al., 2021; Hibberd et al., 2013) and indicate a strong effect of dual-task interference.

**Figure 2.**
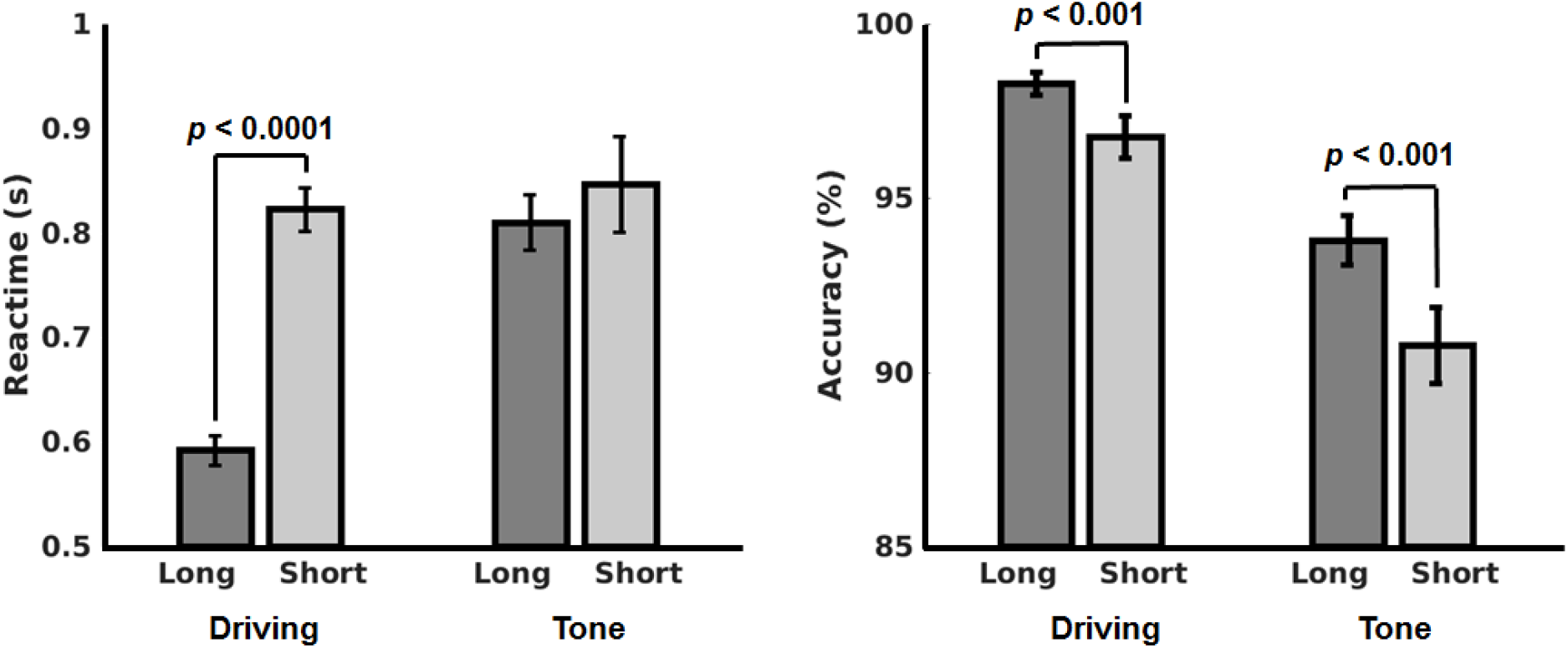
**A)** Reaction times and **B)** Accuracies for the short SOA (dark gray bars) and long SOA (light gray bars) conditions in the driving and the tone tasks. Barplots and errorbars show the mean and standard error, respectively, for each condition.

### FMRI Results

We followed a leave-one-subject-out approach and a union analysis to identify regions that carry information about the driving task in short and long SOA conditions. We identified clusters with above chance decoding accuracy for the driving direction in either the short or the long SOA conditions for the FIR time bin 3 (6 seconds after the stimulus onset) in all but one participant. The mean classification accuracy of these selected ROIs was then compared for two SOA conditions in FIR time bins 2 to 5 in the left-out participant. This procedure was then repeated for all participants. Figure 3A shows a frequency map indicating an overlay of the ROIs of all participants. This figure shows four main regions: the left and right visual cortex, right superior parietal lobe (SPL), and right motor cortex. First, we ran a two-way repeated-measure ANOVA with SOA and FIR time bins as two factors to compare the accuracy for each ROI in the two SOAs across FIR time bins 2 to 5 (Figure 3B). The details of the statistical tests can be seen in table 1. The main effect of SOA was not significant in any of the ROIs (*ps* > 0.05). The interaction of SOA and time bins was significant for the right SPL (*F*(3,57) = 9.32, *p* < 0.0001, two-way repeated measures ANOVA).

**Figure 3.**
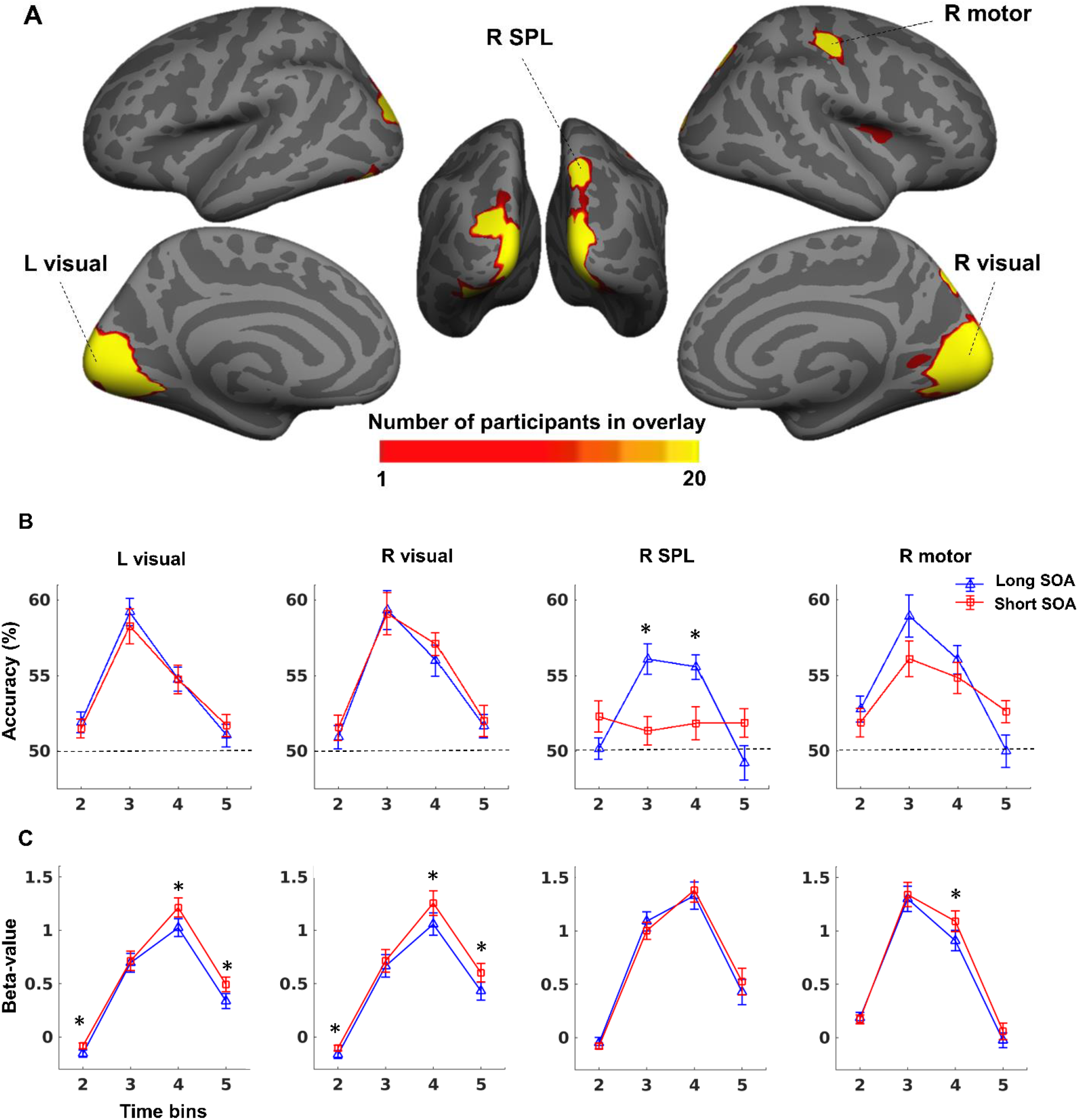
**A)** Frequency maps showing the overlay of ROIs for all participants (n = 20) for the decoding of driving direction. The vertices in the maps are colored based on the number of participants for whom a vertex was included in ROIs. Dashed lines connect the subplots to corresponding ROIs. **B)** Subplots show the mean driving direction decoding accuracy for FIR time bins 2-5 for short (red) and long (blue) SOA conditions in each ROI. Dashed lines show chance-level decoding accuracy (50%). Stars indicate the FIR time bins in which accuracy significantly differed across the short and long SOA conditions (*p* < 0.05, corrected for multiple comparisons). Abbreviations: L = Left hemisphere, R = Right hemisphere.

**Table 1.** Two-way repeated-measure ANOVA for the effect of SOA and time on each ROI’s accuracy. BA denotes the Brodmann area, and VN denotes the number of vertices in each ROI. All p-values were corrected for multiple comparisons across ROIs using FDR correction at q < 0.05, significant p-values are shown in bold text.

| Region Name | BA | MNI coordinate |  |  | NV | SOA |  | SOA x Time |  |
| --- | --- | --- | --- | --- | --- | --- | --- | --- | --- |
|  |  | x | y | z |  | F | p | F | p |
| Left Visual cortex | 17, 18 | -8 | -83 | 3 | 6663 | 0.147 | 0.704 | 0.649 | 0.782 |
| Right Visual cortex | 17, 18 | 7 | -88 | 1 | 5560 | 0.437 | 0.516 | 0.308 | 0.819 |
| Right Motor cortex | 1, 3 | 26 | -68 | 31 | 1200 | 0.578 | 0.456 | 3.460 | 0.051 |
| Right Superior parietal lobe | 7 | 17 | -8 | -10 | 925 | 1.970 | 0.176 | 9.327 | **< 0.001** |

**Table 2.** Comparison of accuracy with chance level (50%) for SOA conditions and pairwise comparison of accuracies between the short and long SOA conditions for each time bin and each region. All p-values were corrected for multiple comparisons across time bins using FDR at q < 0.05. Significant p-values are shown in bold text.

|  |  | FIR time bins |  |  |  |  |
| --- | --- | --- | --- | --- | --- | --- |
|  |  |  | 2 | 3 | 4 | 5 |
| Left Visual cortex | Long > 0.5 | *t* | 2.770 | 9.998 | 5.972 | 1.347 |
|  |  | *p* | **0.016** | **< 0.001** | **< 0.001** | 0.194 |
|  | Short > 0.5 | *t* | 2.387 | 7.143 | 4.947 | 2.307 |
|  |  | *p* | **0.032** | **< 0.001** | **< 0.001** | **0.032** |
|  | Long > Short | *t* | 0.552 | 1.647 | 0.006 | -0.635 |
|  |  | *p* | 0.783 | 0.464 | 0.995 | 0.783 |
| Right Visual cortex | Long > 0.5 | *t* | 1.168 | 7.188 | 5.754 | 2.092 |
|  |  | *p* | 0.257 | **< 0.001** | **< 0.001** | 0.067 |
|  | Short > 0.5 | *t* | 1.855 | 6.552 | 9.503 | 1.937 |
|  |  | *p* | 0.079 | **< 0.001** | **< 0.001** | 0.079 |
|  | Long > Short | *t* | -0.545 | 0.295 | -1.004 | -0.275 |
|  |  | *p* | 0.786 | 0.786 | 0.786 | 0.786 |
| Right Motor cortex | Long > 0.5 | *t* | 3.211 | 6.451 | 6.310 | -0.023 |
|  |  | *p* | **0.006** | **< 0.001** | **< 0.001** | 0.982 |
|  | Short > 0.5 | *t* | 1.994 | 5.194 | 4.732 | 3.596 |
|  |  | *p* | 0.061 | **< 0.001** | **< 0.001** | **0.003** |
|  | Long > Short | *t* | 0.900 | 1.841 | 0.901 | -1.981 |
|  |  | *p* | 0.379 | 0.162 | 0.379 | 0.162 |
| Right Superior parietal lobe | Long > 0.5 | *t* | 0.209 | 6.001 | 6.796 | -0.702 |
|  |  | *p* | 0.837 | **< 0.001** | **< 0.001** | 0.655 |
|  | Short > 0.5 | *t* | 2.216 | 1.401 | 1.651 | 1.959 |
|  |  | *p* | 0.130 | 0.177 | 0.154 | 0.130 |
|  | Long > Short | *t* | -1.718 | 3.935 | 3.024 | -1.872 |
|  |  | *p* | 0.102 | **0.004** | **0.014** | 0.102 |

**Table 3.** Two-way repeated-measure ANOVA for the effect of SOA and time on mean activity (beta-values) in each ROI. BA denotes the Brodmann area, and VN denotes the number of vertices in each ROI. All p-values are corrected for multiple comparisons across ROIs using the false discovery rate of 0.05.

| Region Name | BA | MNI coordinate |  |  | NV | SOA |  | SOA x Time |  |
| --- | --- | --- | --- | --- | --- | --- | --- | --- | --- |
|  |  | x | y | z |  | F | p | F | p |
| **Left Visual cortex** | 17, 18 | -8 | -83 | 3 | 6663 | 24.01 | < 0.001 | 13.62 | < 0.001 |
| **Right Visual cortex** | 17, 18 | 7 | -88 | 1 | 5560 | 26.09 | < 0.001 | 10.86 | < 0.001 |
| **Right Motor cortex** | 1, 3 | 26 | -68 | 31 | 1200 | 7.24 | **0.01** | 11.38 | < 0.001 |
| **Right Superior parietal lobe** | 7 | 17 | -8 | -10 | 925 | 0.77 | 0.78 | 10.20 | < 0.001 |

Focusing on individual time bins in the SPL region, we compared the accuracy for each time bin in each condition to the chance level. The accuracy for decoding the driving direction was above chance level in the third- and fourth-time bins in the long SOA condition (*ts* > 6.01, *ps* < 0.001, paired t-test) and was not significantly greater from chance in any of the time bins in the short SOA condition (*ps* > 0.05). Further pairwise comparisons between short and long SOA conditions in each time bin showed that the decoding of the driving direction for long SOA was significantly higher than the short SOA in time bins third and fourth for SPL (*ts* > 3.02, *ps* < 0.014, paired t-test, see table 4). These results show that although the amount of information about the driving direction does not change in the visual and motor regions, it decreases significantly in the SPL region.

**Table 4.**
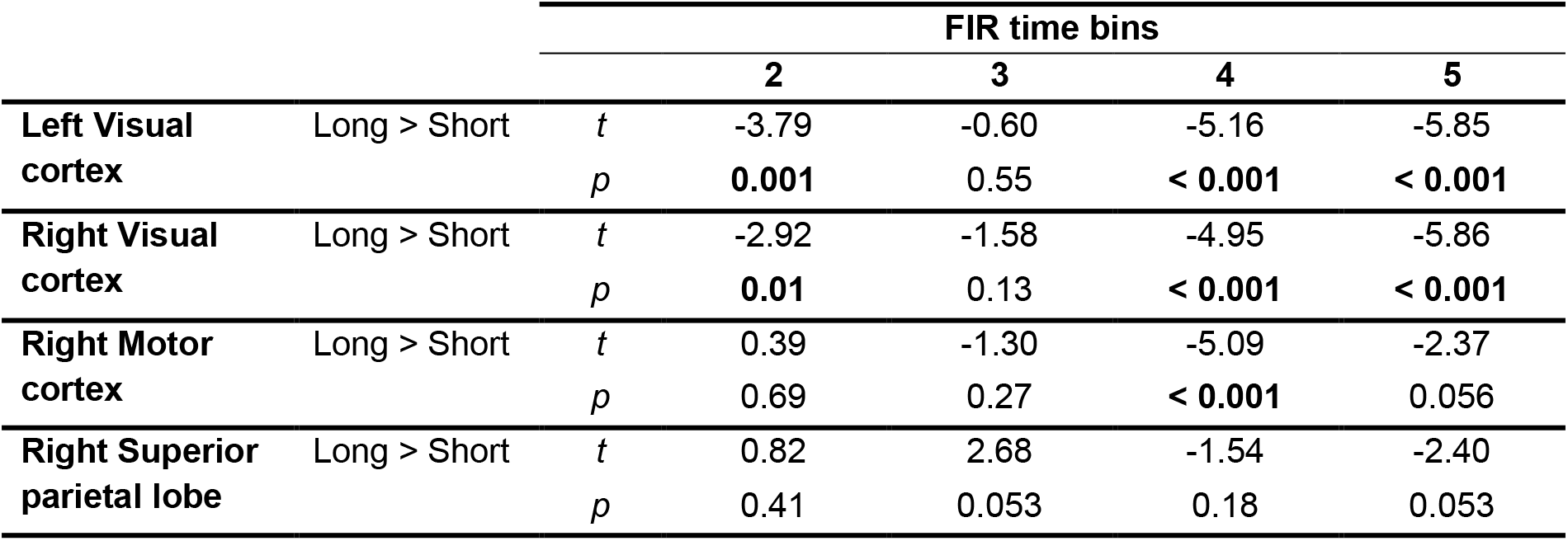
Comparison of mean activity (Beta-values) between the short and long SOA conditions for each time point and each region. All p-values are corrected for multiple comparisons across time points using FDR 0.05.

We also examined the mean activity in regions with information about the driving task (Figure 3C). For this purpose, we ran a two-way repeated-measures ANOVA with SOA and FIR time bins as two factors. The details of the statistical tests can be seen in table 3. Results indicated the main effect of SOA was significant for both right and left visual and motor regions (ps < 0.05,), but it was not significant for the SPL (p = 0.78). However, the interaction of SOA and time was significant for all regions (ps < 0.05).

To further investigate the effect of SOA across FIR time bins, we ran a paired t-test for each time bin in each region. The statistical details can be seen in table 4. For the left and right visual regions, the mean activity in the short SOA was significantly higher than the long SOA in the time bins 2, 4, and 5. For the right motor cortex, only mean activity in the time bin 4 was significantly higher in short SOA compared to the long SOA and in other time bins, the effect of SOA was not significant. However, for the right SPL in all time bins, the SOA effect was not significant. The comparison of the results of mean activity and classification accuracy in the informative driving regions revealed, although the mean activity increased in the visual and motor regions in the short SOA compared to the long SOA condition, the driving information did not change in these regions. In contrast, the mean activity did not change in the SPL region across SOA and FIR time bins, but the driving information decreased across time in the short SOA versus long SOA.

We next investigated if the decoding accuracies observed in our ROIs are related to behavior. A Pearson correlation analysis revealed a negative correlation between the turn direction decoding accuracy in time bin 3 in the SPL region and the reaction time of the driving task obtained from the behavioral results (*r* = −0.507, *p* = 0.032, corrected for multiple comparisons, Figure 4). This correlation was not significant for the other three regions for all FIR time bins. Moreover, there was no correlation between mean activity and the driving reaction time for all regions (*ps* > 0.05). These results suggest that the drop of information in the SPL region could play a role in dual-task interference and the drop in performance when the two tasks are performed concurrently.

**Figure 4.**
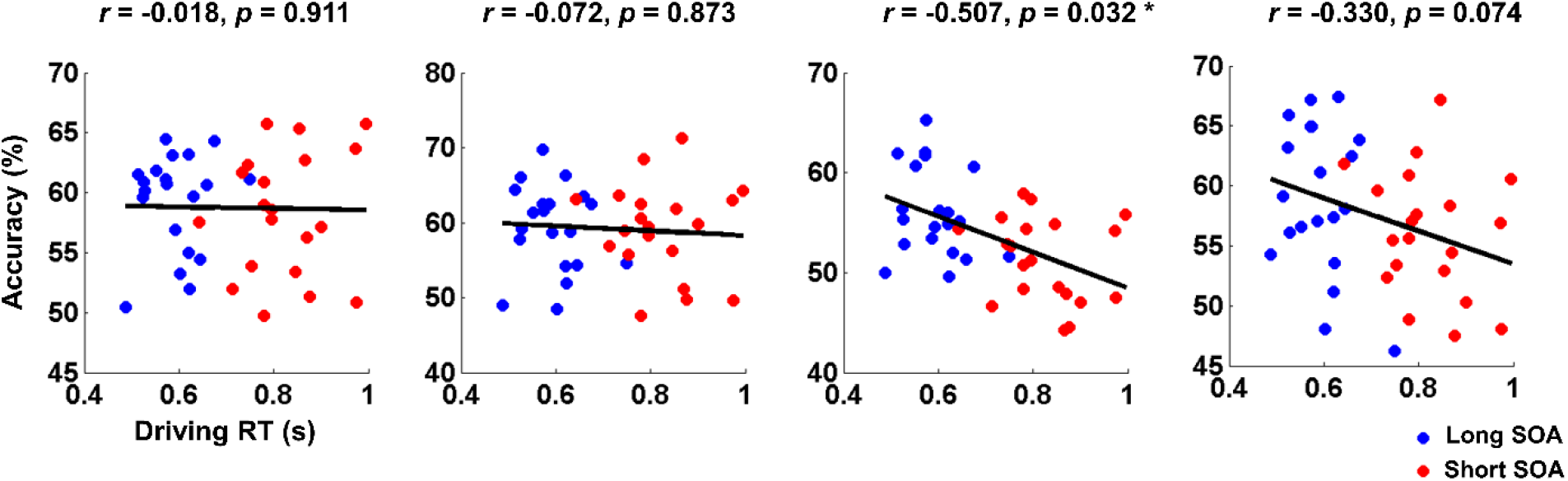
The Pearson correlation between the decoding accuracy of the driving direction in FIR time bin 3 and behavioral reaction time of driving direction for ROIs. Abbreviations: L = Left hemisphere, R = Right hemisphere.

## Discussion

Here, we investigated the effect of dual-task interference on the BOLD response in a dual-task paradigm in which participants performed a driving turn in a simulated driving environment along with a tone discrimination task. The two tasks were either presented close together (short SOA) or far from each other (long SOA) in time. The behavioral results showed an increase in RT and a decrease in the accuracy of the driving task in the short SOA compared to the long SOA trials consistent with previous studies (Hibberd et al., 2013; Levy et al., 2006), confirming that the performance is influenced by dual-task interference. We performed a time-resolved fMRI analysis by investigating the patterns of brain activity at multiple time bins after the stimulus onset. The results of MVPA revealed regions that carry information about the driving direction including the superior parietal lobe (SPL), visual, and motor regions. Comparing the decoding accuracies across short and long SOA conditions, we showed for the first time that the information for the driving direction gets disrupted by dual-task interference in the right SPL but not in the visual and motor regions.

Although the driving information decreased in the SPL region in the short SOA compared to the long SOA, the mean activity did not vary in this region in all FIR time bins across SOA conditions. This finding indicates that the dual-task interference disrupts the content of driving information in the SPL without changing the average activity of this region. Surprisingly, the behavior-brain correlations also showed a negative correlation between the reaction time of the driving turn and the driving information in the SPL region. These results suggest that disruption in the driving information in the SPL is likely to play a role in driving performance decline. To the best of our knowledge, this finding is novel and has not been reported in previous studies.

SPL has been shown to play a role in the maintenance of object shape and location in working memory (Rowe, Toni, Josephs, Frackowiak, & Passingham, 2000; Xu & Chun, 2006), maintenance and manipulation of auditory stimuli (Koenigs, Barbey, Postle, & Grafman, 2009) and contain more information about an attended compared to an unattended item (Vaziri-Pashkam & Xu, 2017).

For our driving task, it is possible that the SPL would be involved in maintaining the information about the driving task in working memory until the processing of the tone task finishes. In the short SOA condition, because the driving trigger was presented immediately after the tone task, the maintenance of information for the driving task could be disrupted by the tone task. This could explain the decrease in the decoding of driving turn information in the SPL in the short SOA and a delay in discriminating the driving turn direction by the participants.

SPL receives input directly from the visual cortex (Caspers & Zilles, 2018; Xu, 2018). Our univariate analysis revealed clear signs of the change in the shape of HRF in the visual cortex during dual-task conditions. Nevertheless, the information for the driving direction was not significantly affected in the visual cortex. The increase of activity of the visual region across FIR time bins in the short compared to the long SOA might be due to top-down feedback signals to sensory regions to increase the activity in these regions or keep them active for longer durations to compensate for the disruption in the driving information caused by dual-task interference.

Our results also indicated a change in the shape of HRF in motor regions with an increase in the response in later time points, without any change in the information content for driving direction. Similar to the increase in signal in the visual regions, this change might be related to the increase of the signal to preserve the information contained in the motor regions during dual-task interference. They may also be related to possible delays in the processing of the motor stage during dual-task interference that resulted from the interference in the previous stage of processing. These results are novel and to the best of our knowledge have not been reported in previous fMRI studies of dual-task interference. The delays in the motor stage of processing have been proposed in previous behavioral modeling studies (Zylberberg et al., 2012) and are in line with our previous behavioral and modeling findings (Abbas-Zadeh et al., 2021).

A recent Magnetoencephalography (MEG) study by Marti et al. (2015), has shown that the information is disturbed during the central processing stage of the task (about 350-450 ms after the onset of the stimulus, where the decision is being made and the response is being selected). However, due to limitations in the spatial resolution of the MEG signal, it is not possible to attribute the observed information disturbance to a particular location in the brain. The findings of the current study propose SPL as a candidate region in which the disturbance of information leads to dual-task interference. Needless to say, the causal role of SPL in dual-task performance can only be established using future studies with causal methods that allow for manipulation of the signal in SPL.

In sum, we show an association between the information content of SPL and dual-task performance in a simulated driving environment. These results extended our understanding of the neural correlates of dual-task interference and are informative for formulating biologically plausible models of dual-task interference that apply to more everyday settings.

## Notes

### Competing Interest Statement

The authors have declared no competing interest.

### Summary of Updates

The result of the univariate whole-brain analysis was removed from the paper because it was confusing. Therefore, we had to update all sections of the manuscript.

